# KiPERT predicts responses to single- and double-gene perturbations and transfers perturbation effects across cell lines

**DOI:** 10.64898/2026.11.19.767987

**Authors:** Qiantong Liang, Yi Wang, Qing Rex Lyu

## Abstract

Distinguishing transcriptional responses to genetic perturbations is essential for understanding gene function. However, models may reproduce shared expression changes while missing perturbation-specific and context-dependent responses. Here we present KiPERT (Knowledge-informed Perturbation Effect Regression and Transfer), which combines biological similarity kernels with kernel regression to predict mean expression changes for unseen single-and double-gene perturbations. For prediction across cell lines, KiPERT transforms perturbation effects in training cell lines using optimal transport between control cell distributions, requiring only controls from the new cell line. Across evaluated Perturb-seq datasets, KiPERT improved average performance over tested methods in perturbation discrimination, prediction of response direction and identification of differentially expressed genes. In an in vivo mouse cardiac reprogramming dataset, KiPERT reduced mean squared error in cardiac transcriptional response estimates by 40% relative to sample means and captured the known upregulation of cardiac genes following *Calr* knockdown across batches.

## Introduction

Artificial intelligence virtual cells (AIVCs) aim to represent cellular states across molecular modalities and predict how those states change under different conditions and perturbations^1,2^. One task within this broader effort is to predict how genetic perturbations alter gene expression in each cellular context^1,3,4^. Perturb-seq combines pooled genetic perturbations using sgRNAs and single-cell RNA sequencing (scRNA-seq), providing ideal training data by linking individual genetic perturbations to corresponding transcriptome changes at scale^5–8^. Perturb-seq profiles transcriptional responses to a defined set of perturbations in the cells studied; however, responses to genetic and chemical perturbations can depend on cellular context^9,10^. Thus, two key challenges for AIVCs are: (1) predicting responses to unseen perturbations within a given cellular context and (2) inferring unmeasured responses in a target cellular context from responses to the same perturbations measured in a source context^3,4^.

Several computational methods predict transcriptional responses to unseen genetic perturbations within a given cellular context^11^. Graph-based methods such as GEARS and TxPert use prior knowledge of gene-gene relationships to extrapolate from measured responses to held-out single-gene perturbations and gene combinations^3,12^. Scouter combines embeddings derived from textual gene descriptions with control-cell expression profiles to predict responses to unseen single- and double-gene perturbations^13^. Single-cell foundation models such as scGPT and scFoundation leverage representations learned from large-scale transcriptomic datasets to predict post-perturbation expression profiles^14,15^. Conditional generative models such as PerturbNet use gene functional annotations to predict the distribution of single-cell expression states following unseen CRISPR perturbations^16^. Despite these advances, existing models can achieve high agreement with measured expression profiles by capturing shifts shared across perturbations, yet still struggle to recover the transcriptional changes that distinguish individual unseen single- and double-gene perturbations^17^.

Available Perturb-seq datasets cover cellular contexts unevenly, with several foundational large-scale screens conducted in established cell lines such as K562 and RPE1^18^. Extending pooled CRISPR screening to primary cells can be limited due to difficulties in maintaining cell states ex vivo and inefficient lentiviral delivery of the Cas9 enzyme^19,20^. Although genome-scale Perturb-seq has now been demonstrated in primary human CD4□ T cells, establishing stable lentiviral expression of genome-editing enzymes remains challenging in primary human hematopoietic stem and progenitor cells^21,22^. Two complementary approaches can help address these gaps. In vivo Perturb-seq enables direct profiling of transcriptional responses within tissues with heterogeneous cell types and has progressed from targeted screens in the developing mouse brain to large-scale screens across major brain regions^23–25^. Computational transfer uses responses measured in a source context to infer the effects of the same perturbations in a target context where their responses have yet to be measured^26^.

Here we introduce KiPERT (Knowledge-informed Perturbation Effect Regression and Transfer), which combines biological priors with measured responses through kernel regression to predict mean expression changes for unseen single- and double-gene perturbations and model non-additive effects. For predictions across cell lines, KiPERT uses optimal transport between control-cell expression distributions to map the effects of the same perturbation to a target line. Across the evaluated datasets, KiPERT improved average performance in perturbation discrimination and response direction prediction, including transfer across four cell lines. We further applied KiPERT to an in vivo cardiac reprogramming dataset to improve response estimates for sparsely sampled perturbations and capture known cardiac transcriptional responses.

## Results

### Overview of KiPERT

KiPERT predicts perturbation effects in mean gene expression relative to control cells. Within a cellular context, it predicts responses to single- and double-gene perturbations not seen in training. For prediction across cell lines, the query perturbation must have been measured in at least one training cell line, while only control cells are required from the target cell line.

Biological prior knowledge is encoded as gene similarity kernels derived from 12 biological priors, including genetic dependencies, gene representations, protein interactions, pathways, regulatory relationships, cellular localization, and functional annotations (**Fig. 1a**). The kernels can be combined uniformly or with optional weighting based on alignment with training responses. A mixing coefficient *β* controls the alignment contribution, with *β* = *0* giving uniform weights. The main single- and double-gene benchmarks in this study used *β* = 0.5; alternative values were evaluated in sensitivity analyses. Kernel ridge regression uses the resulting kernel to predict mean expression changes.

**Fig. 1.**
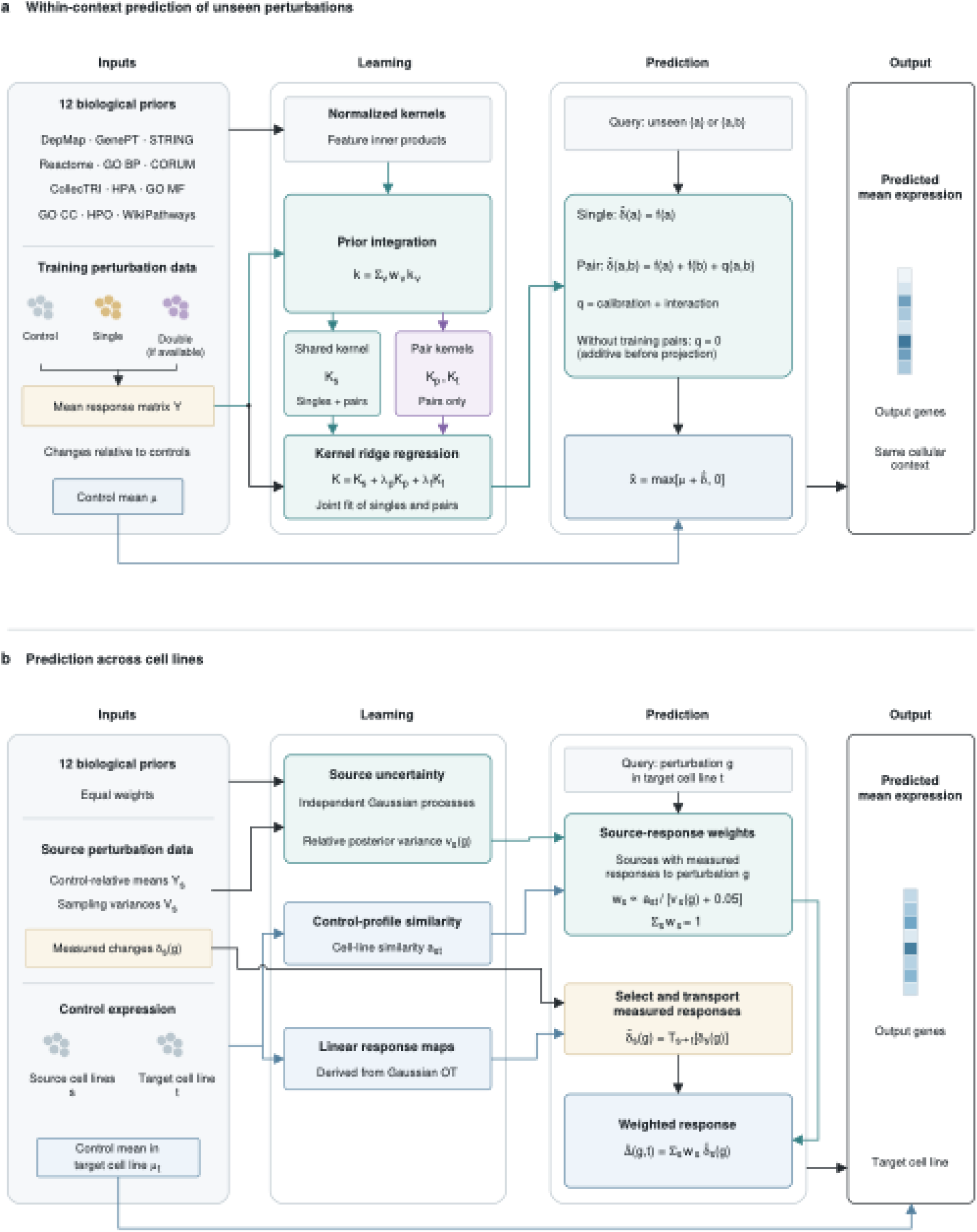
Overview of KiPERT. **a,** Prediction of responses to unseen single- and double-gene perturbations within a cellular context. **b,** Transfer of measured perturbation responses across cell lines. GP, Gaussian process; OT, optimal transport.

For double-gene perturbations, single- and double-gene perturbations are fitted jointly using a shared first-order kernel and pair-specific calibration and interaction kernels. The predicted change for a gene pair is 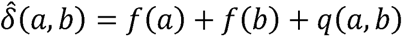, where the shared components *f*(*a*) and *f*(*b*) are estimated jointly from single- and double-perturbation data, and *q*(*a*, *b*) denotes the pair-specific contribution. Without training combinations, *q*(*a*, *b*) = *0* and the response is additive before non-negative expression projection. Predicted changes are added to the control mean, and negative expression estimates are set to zero.

For prediction across cell lines, KiPERT uses an equally weighted combination of the 12 kernels and fits a Gaussian process independently in each training cell line (**Fig. 1b**)^27^. Control-cell profiles define similarities between cell lines and linear transformations derived from Gaussian optimal transport (OT)^28^. Measured responses to the query perturbation are transformed and combined using weights based on control-cell similarity and relative posterior variance. Only source lines with measurements of that perturbation contribute. The transferred response is then added to the target control mean. Transfer can also use an identity map, partial transport or uniform source weights. The main benchmarks here used full OT.

### Single-gene perturbations

We evaluated unseen single-gene perturbations across five datasets spanning iPSC-derived neurons (Tian CRISPRi and CRISPRa), K562 cells (Adamson and Norman singles), and melanoma cells (Frangieh). Comparisons with biolord, Scouter, TxPert, GEARS, STAMP-Geneformer, and RegFormer-GEARS used three computational splits. Across the ten metrics in **Fig. 2a**, KiPERT achieved the best or joint-best average rank on nine metrics, with each dataset weighted equally. Its average rank was 1.4 for both top-100 expression mean squared error (MSE) and differentially expressed gene (DEG) area under the precision–recall curve (AUPRC), 1.5 for precision and recall at 100, and 1.6 for response-sign agreement. KiPERT and biolord shared the best average rank of 1.8 for all-gene MSE. These comparisons support improved average performance in expression prediction and DEG recovery among the evaluated methods. After subtracting the shared transcriptional response from both predicted and observed expression changes, KiPERT achieved a mean Pearson correlation of 0.472 in Adamson, compared with 0.304 for GEARS, the next-highest method (**Fig. 2b**). The shared response was estimated as the average training response weighted by cell count.

**Fig. 2.**
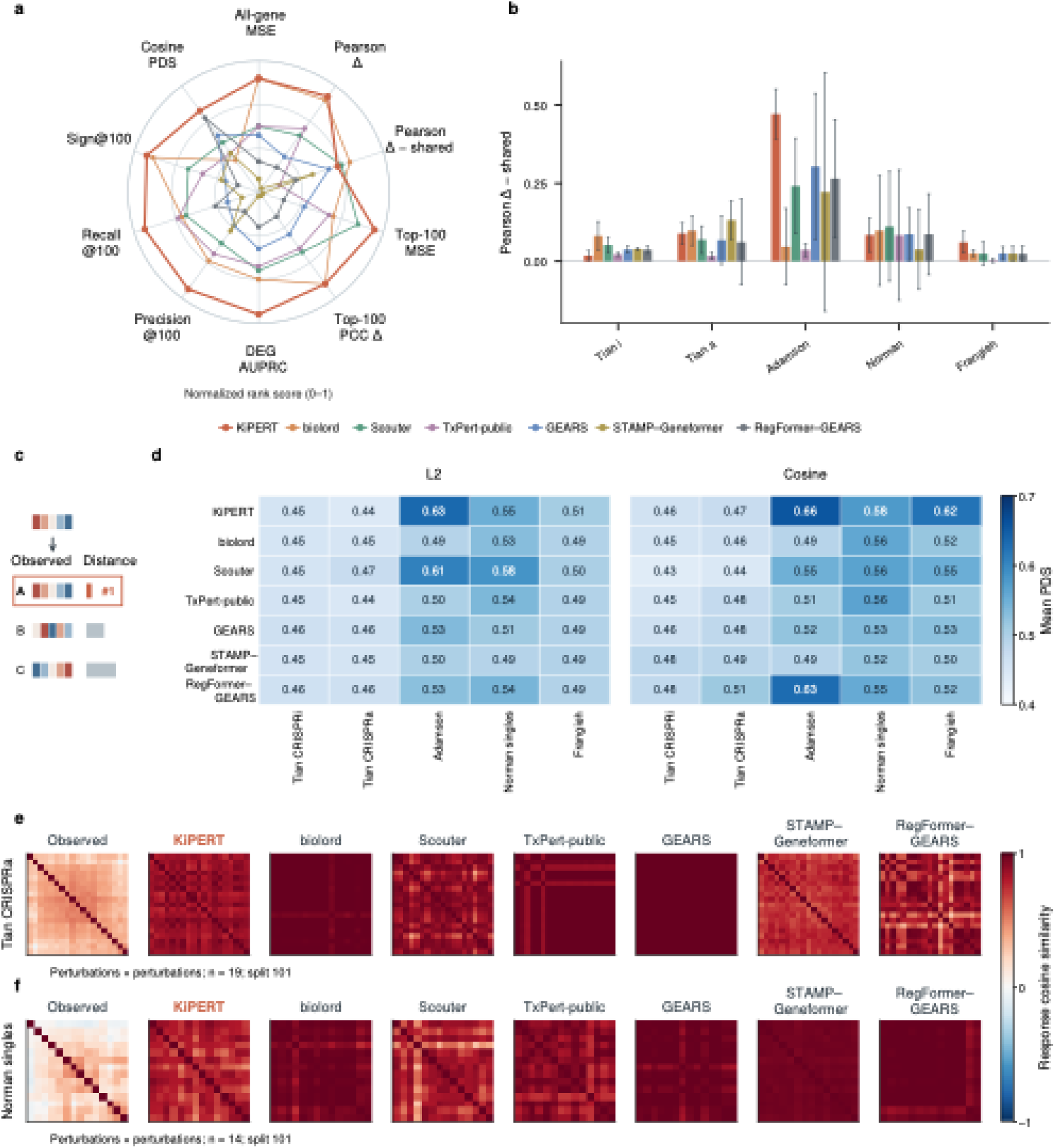
Prediction of unseen single-gene perturbations. **a**, Performance ranks for seven methods on ten metrics. Larger values indicate better ranks. Δ denotes the difference in mean expression between perturbed and control cells. Top-100 MSE, PCC and Sign@100 use genes ranked by absolute observed Welch t-statistic; Sign@100 measures response-direction agreement. **b**, Pearson correlation after subtracting the shared training response from observed and predicted changes. Bars show means and error bars show 95% Student-t confidence intervals across three perturbation splits. **c**, Illustration of perturbation discrimination. **d**, Mean perturbation discrimination scores (PDS) for L2 and cosine distances. Higher scores indicate better identification. **e,f**, Pairwise cosine similarities of observed and predicted response profiles for Tian CRISPRa and Norman singles. Blue, white and red indicate negative, zero and positive similarity. MSE, mean squared error; PCC, Pearson correlation coefficient; DEG, differentially expressed gene; AUPRC, area under the precision–recall curve.

Perturbation-specific predictions were further assessed by comparing each predicted profile with observed responses from the same context and split, excluding the perturbed gene from the distance calculation (**Fig. 2c**). The perturbation discrimination score (PDS) quantifies the rank of the observed response among the candidates. KiPERT achieved the highest cosine PDS in Adamson, Norman singles and Frangieh, with scores of 0.656, 0.576 and 0.622, compared with 0.631, 0.565 and 0.550 for the best corresponding comparators (**Fig. 2d**). In the illustrated Tian CRISPRa and Norman singles contexts, pairwise response-similarity matrices showed variation across KiPERT prediction responses (**Fig. 2e,f)**.

### Prior information

To determine whether specified gene relationships improved perturbation prediction, we reassigned prior information to different genes, creating a control with mismatched gene identities and prior information. With equal weighting of the five datasets, cosine PDS increased from 0.5002 for the control to 0.5565 for the model. This corresponded to an 11.26% relative increase (95% paired bootstrap interval, 8.00–14.69%). DEG AUPRC increased from 0.1253 to 0.1601, a 27.72% relative improvement (18.51–39.38%). The results therefore support the predictive value of linking biological prior information to gene identities (Extended Data Fig. 1a,b).

Expanding the prior set improved average prediction, with diminishing additional gains as the set approached all 12 priors. Relative to the mean performance of 2-prior subsets, the full model improved cosine PDS by 5.33% and DEG AUPRC by 21.00% (Extended Data Fig. 1c,d). We assessed the contribution of each of the 12 priors by removing one view at a time. Removing DepMap reduced cosine PDS by 2.21% (95% paired bootstrap interval, 0.73–3.78%), mainly in Frangieh; most other individual removals produced relatively smaller PDS changes. Compared with a 3-prior model comprising DepMap, GenePT and STRING, the full model increased DEG AUPRC by 16.88% (10.24–25.52%) (Extended Data Fig. 2).

Prior groups contributed differently across prediction objectives. The high-coverage group comprised DepMap, GenePT, STRING and the Human Protein Atlas (HPA). The full model increased DEG AUPRC by 22.22% relative to the four-prior group (95% paired bootstrap interval, 14.71–31.88%). Stratifying targets by prior similarity to training genes gave relative AUPRC improvements over the jointly permuted control of 16.43%, 31.54% and 34.96% in the low-, intermediate- and high-proximity groups (Extended Data Fig. 3). Besides, optional alignment weighting provided a modest benefit for DEG recovery under a common scoring procedure (Extended Data Fig. 4).

### Double-gene perturbations

We evaluated held-out gene combinations in Norman, Replogle exp6, Wessels L4 and Dixit K562 under three levels of single-perturbation availability. Seen 0, seen 1 and seen 2 provided measurements for zero, one or both constituent single-gene perturbations, while retaining the same other training combinations. Across five computational splits, KiPERT had the lowest mean opposite-direction fraction on the 20 genes with the largest absolute observed expression changes among the methods in **Fig. 3a–c**. With equal weighting of the four datasets, these fractions were 0.081, 0.069 and 0.059 for seen 0, seen 1 and seen 2, compared with 0.099, 0.095 and 0.084 for the best displayed comparators. Increasing the availability of single-perturbation measurements also reduced top-20 normalized MSE, from 0.250 to 0.177 and 0.110 in Norman, and from 0.232 to 0.206 and 0.189 in Replogle exp6 (**Fig. 3d,e**).

**Fig. 3.**
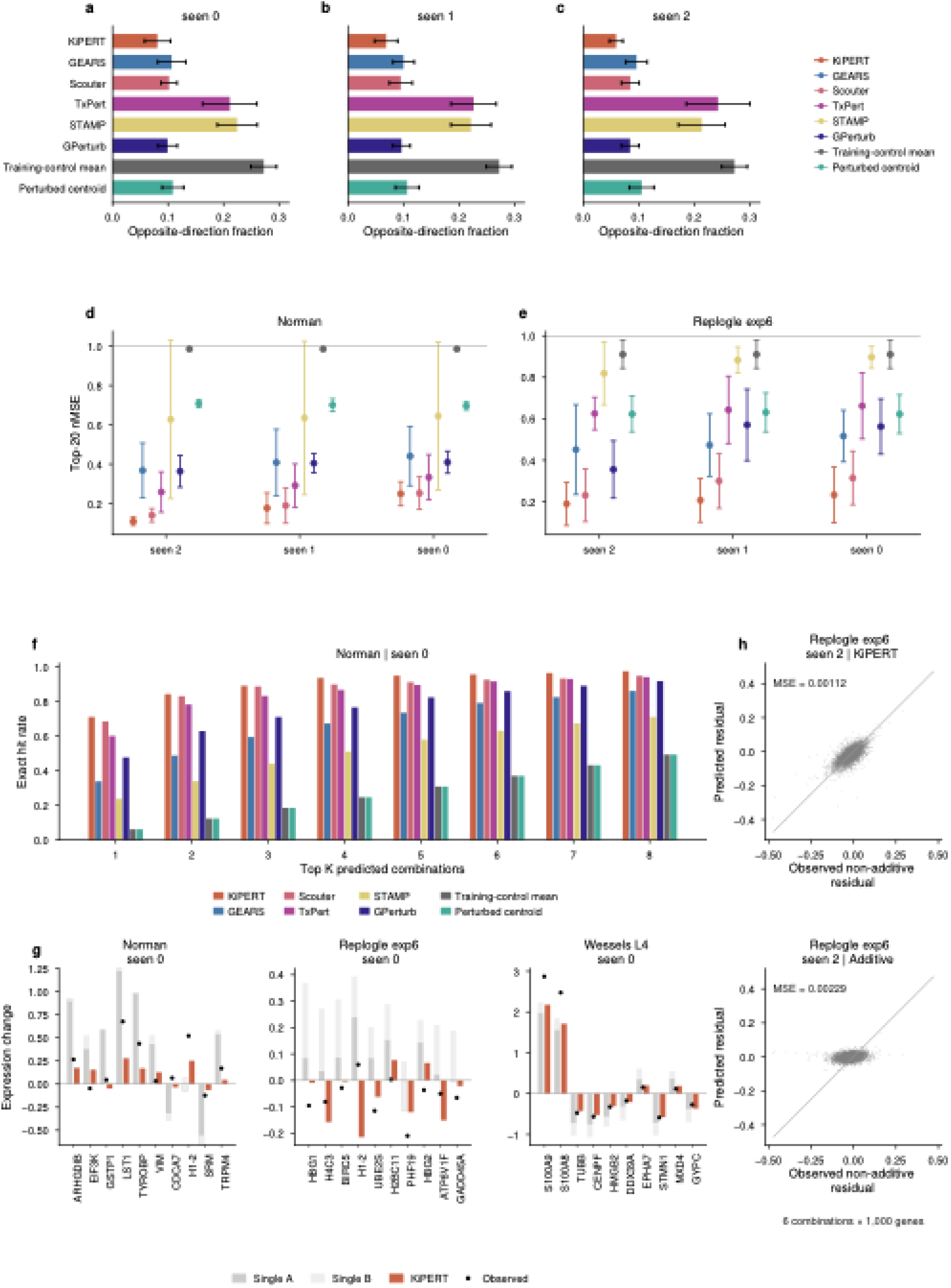
Prediction of unseen double perturbations. Seen 0, seen 1 and seen 2 provide zero, one or both single perturbations during training, respectively. **a–c**, Opposite-direction fractions for seen 0–2 among the 20 genes with the largest absolute observed expression changes, averaged equally across Norman, Replogle exp6, Wessels L4 and Dixit K562. **d,e,** Top-20 normalized mean squared error (nMSE) in Norman and Replogle exp6, relative to the mean squared observed change. Lower values indicate better predictions. **f,** Fraction of observed combinations recovered among the top K predicted combinations ranked by Pearson similarity, in Norman seen 0. **g,** Expression-change examples from seen 0 (*SPI1* + *MAP2K6* in Norman, *HUS1* + *SREBF2* in Replogle exp6 and *GSE1* + *INTS1* in Wessels L4). Each displays the ten genes with the largest absolute departure from the sum of two single perturbations. Grey segments show separately single effects. **h,** Observed versus predicted residuals for KiPERT (top) and the additive predictor (bottom), in Replogle exp6 seen 2. All six combinations × 1,000 genes are shown. Grey diagonals indicate equality.

For each observed response, we ranked the predicted profiles of held-out combinations by Pearson similarity. In Norman seen 0, KiPERT ranked the correct combination first for 71.0% of queries, compared with 68.4% for Scouter, 60.0% for TxPert and 33.8% for GEARS (**Fig. 3f**). The correct combination appeared among the top four and top eight candidates in 93.5% and 97.4% of queries, respectively.

The expression examples illustrated predicted departures from the sum of measured single-perturbation effects (**Fig. 3g**). For *ARHGDIB* in the Norman example, the constituent effects summed to 0.894, whereas the observed double-perturbation change was 0.266 and KiPERT predicted 0.175. We quantified these departures in the Replogle exp6 seen-2 split shown in **Fig. 3h** using the contrast *R* = *Y*_*AB*_ *— A*_*train*_ *— B*_*train*_ *+ c_score_*. The same measured single-perturbation reference and independent scoring-control mean were used for observed and predicted combination profiles. Across six held-out combinations and 1,000 genes, KiPERT achieved a contrast MSE of 0.00112, compared with 0.00229 for the available-evidence additive predictor, a reduction of 51.0%. The additive predictor summed single-perturbation effects relative to training controls and applied non-negative expression projection. KiPERT therefore predicted the expression changes in these held-out combinations more accurately than this additive baseline.

Prior contributions to double-perturbation prediction depended on the available single-perturbation measurements. Removing CORUM increased mean top-20 normalized MSE relative to the full model by 3.35%, 2.95% and 1.74% in seen 0, seen 1 and seen 2, respectively. These increases support a contribution from protein-complex information (Extended Data Fig. 5).

### Prediction across cell lines

We evaluated prediction across cell lines by withholding perturbation profiles from K562, HepG2, RPE1 and Jurkat in turn. Each target cell line contributed control cells, and every test perturbation was measured in at least one training cell line. The benchmark comprised 3,712 cell-line–perturbation conditions across 6,642 genes, representing 1,595 distinct perturbations: 884 conditions in K562, 867 in HepG2, 983 in RPE1 and 978 in Jurkat.

Across the five methods shown in **Fig. 4a–e**, KiPERT achieved the highest mean score on all five metrics, with equal weighting of the four cell lines. These metrics assessed response-direction agreement, perturbation discrimination, expression-change correlation, log-fold-change correlation among observed differentially expressed genes (DEGs), and DEG overlap. The DEG-overlap score was 0.217, compared with 0.134 for TxPert and 0.066 for STATE. The *SEH1L* example illustrated agreement in mean expression across genes, with squared Pearson correlations of 0.972–0.983 across the four cell lines (**Fig. 4f**).

**Fig. 4.**
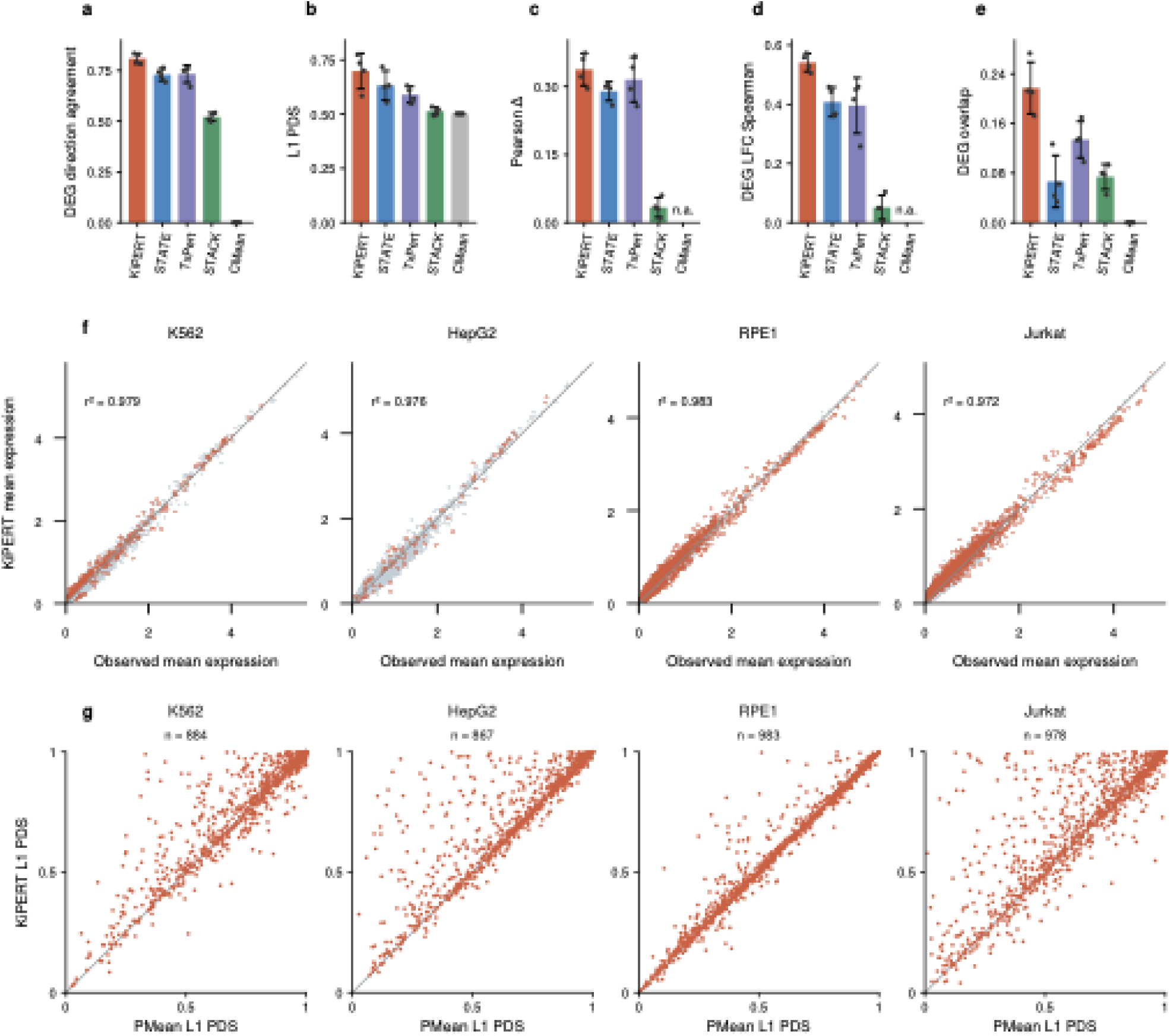
Transfer of perturbation responses across cell lines. **a–e**, Response-direction agreement among observed differentially expressed genes (DEGs; **a**), L1 perturbation discrimination score (PDS; **b**), Pearson correlation of expression changes (**c**), Spearman correlation of log-fold changes among observed DEGs (**d**), and DEG overlap (**e**). Higher values indicate better performance. Bars show equally weighted means across four cell lines, black points show cell-line scores and error bars show sample standard deviations. n.a. denotes undefined correlations. DEGs have adjusted *P* < 0.05. For overlap, predicted significant genes are ranked by absolute log-fold change and compared with observed DEGs at the observed DEG count. **f**, Observed versus KiPERT-predicted mean expression after *SEH1L* perturbation. Points represent genes; orange marks observed DEGs and grey marks other genes. The displayed *r*² is squared Pearson correlation. **g**, Paired L1 PDS for PMean (horizontal) and KiPERT (vertical). Dashed diagonals in **f,g** indicate equality.

KiPERT also improved response discrimination relative to PMean, which averages measured responses to the same perturbation across available source cell lines (**Fig. 4g**). Mean L1 perturbation discrimination scores were higher in all four target cell lines, with higher scores for 57.5–63.7% of conditions within each cell line. The equally weighted mean increased from 0.660 to 0.699, a relative increase of 5.88% (95% perturbation-cluster bootstrap interval, 5.25–6.57%). All-gene expression-change MSE, averaged equally across cell lines, was higher for KiPERT than PMean with a relative increase of 7.38%, indicating differing performance in perturbation discrimination and gene-wise prediction error.

Full OT increased mean L1 PDS from 0.6639 for the identity map to 0.6985, a relative improvement of 5.22% (95% perturbation-cluster bootstrap interval, 4.68–5.77%). PDS increased in all four target cell lines. The PDS advantage persisted after reselecting the identity-map temperature using source cell lines. Mean PDS increased as the transport-strength *ç* rose from 0 to 1, whereas intermediate transport strengths gave lower MSE. This comparison revealed a trade-off between response discrimination and amplitude error (Extended Data Fig. 6).

### Cardiac reprogramming responses from sparse perturbation measurements

Low cell coverage presents an additional challenge for interpreting perturbation responses in biological screens. We applied KiPERT to a Perturb-seq dataset of in vivo cardiac reprogramming in mice, using cells collected 14 days after myocardial infarction^29^. Within each sample batch, kernel regression combined five or ten input cells per measured perturbation with data from other measured perturbations to estimate its expression profile (**Fig. 5a**). We quantified the cardiac response as the mean expression change across eight cardiac genes relative to non-targeting (NT) controls. The cardiac expression-change MSE decreased from 0.03213 for the sample mean to 0.01927 for KiPERT, a 40.0% reduction (95% perturbation-cluster bootstrap interval, 30.0–46.3%; **Fig. 5b,c**). Mean cardiac-gene expression before NT subtraction also showed lower error, with a reduction of 49.7% (39.2–56.1%). With ten input cells, estimated error reductions were 22.7% for expression change (−0.6 to 36.3%) and 30.1% for mean expression (−4.9 to 46.5%).

**Fig. 5.**
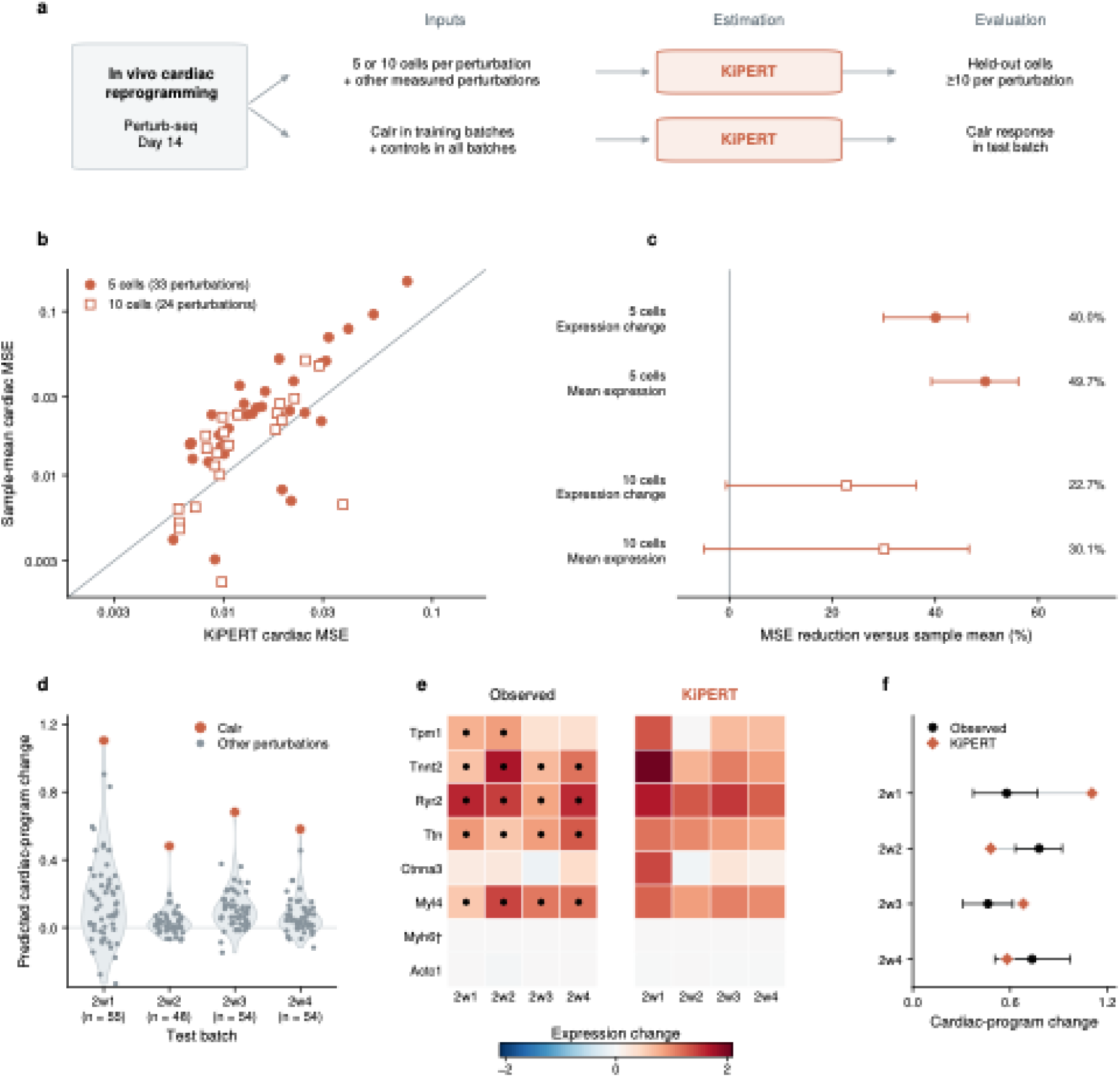
Sparse-measurement estimation and transfer of cardiac perturbation responses. **a,** KiPERT applications to in vivo cardiac reprogramming Perturb-seq at day 14 after myocardial infarction. **b,** Cardiac expression-change mean squared error (MSE) for KiPERT (horizontal) and sample means (vertical), on logarithmic axes. **c,** Relative MSE reduction, 100 × (MSE(sample mean) − MSE(KiPERT))/MSE(sample mean), for expression change and mean expression before NT subtraction. **d,** *Calr* response within all predictions in four test batches. **e,** Observed and predicted *Calr* changes across eight cardiac genes with blue and red indicating negative and positive changes. **f,** Observed cardiac-program changes (black points) and intervals, connected to KiPERT estimates (coral diamonds).

*Calr* knockdown provided a reported positive reference for evaluating transfer of a cardiac response across sample batches. For each test batch, KiPERT transferred measured *Calr* responses from the other batches using Gaussian optimal transport, with NT controls available from all batches. *Calr* ranked first in the four test batches (**Fig. 5d**). The predicted *Calr* response preserved the observed upregulation of *Tnnt2*, *Ryr2*, *Ttn* and *Myl4* (**Fig. 5e,f**).

## Discussion

KiPERT preserved perturbation-specific information across single- and double-gene prediction and transfer between cell lines. The evaluation results distinguish reproducing an expression profile from resolving the effects of individual perturbations. Correlations of expression changes can be driven by shifts shared across perturbations, as Systema highlights.^17^ KiPERT retained predictive information after shared-response subtraction and discriminated among perturbations using predicted response profiles. These findings support the retention of information beyond the common transcriptional component. Direction agreement, response discrimination and DEG recovery assess different properties of that information.

Biological priors provide a statistical structure for sharing responses. GEARS and TxPert also use gene relationships to inform perturbation prediction.^3,12^ KiPERT expresses these relationships as similarity kernels, making the connection between prior information and regression explicit. Its central assumption is that relationships encoded by the priors are informative about response similarity in the cellular context being modelled. Gene permutation with preserved coverage reduced performance, supporting the predictive information in the correspondence between genes and prior representations. Prior-number and group comparisons further indicated that different definitions of relatedness supported complementary prediction objectives, with diminishing gains near the full set. Some compact sets remained competitive for discrimination, whereas the full set provided clearer benefits for DEG recovery. Prior selection therefore depends on both information content and the intended prediction objective. Besides, KiPERT treats alignment as an optional extension of uniform integration. Applications can assess these settings using independent validation data matched to the intended objective.

The cardiac application illustrates how these models can assist interpretation of an existing experiment. KiPERT reduced estimation error relative to sample means in the five-cell analysis, which supports model estimation of cardiac responses from limited numbers of cells. The *Calr* analysis addresses preservation of a reported response.^29^ Positive cardiac-gene changes were retained. Such estimates can support interpretation of low-coverage conditions and comparison of expression patterns across batches.

This study has several limitations. First, KiPERT predicts condition-level mean expression rather than single-cell distributions and cannot distinguish transcriptional changes within a cell state from changes in cell-state composition. Second, its predictions depend on prior coverage and on whether prior similarity reflects response similarity in the context studied. Correlated priors may encode redundant information. Third, transfer across cell lines requires that the query perturbation has been measured in at least one source cell line and that representative target controls are available. The Gaussian control-based map assumes that baseline covariance differences inform perturbation-specific context dependence; it may be insufficient for nonlinear, multimodal or perturbation-induced changes not reflected in controls. Broader prospective validation, including additional primary and in vivo contexts and functional readouts, is needed to establish where these predictions reliably support experimental design.

## Methods

KiPERT predicts changes in mean gene expression following genetic perturbation. Within a cellular context, we withheld single- or double-gene responses from model fitting. For transfer across cell lines, the query perturbation was measured in at least one source cell line and control cells were available in the target cell line. For perturbation *p* in context *c*, let *x̅_p,c_* denote its mean expression vector and *x̅*_0,*c*_ the corresponding control mean. The response vector was *δ_p,c_* = *x̅_p,c_* - *x̅*_0,*c*_. Each vector contained one entry per gene on the evaluation axis. Within-context models were fitted to these condition means. Predicted responses were converted to expression means by 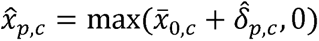 with the maximum applied elementwise. The control partitions and preprocessing used for each benchmark are specified below.

### Biological priors

We represented perturbation targets using 12 sources of biological knowledge: DepMap genetic dependencies, GenePT gene embeddings, STRING protein network embeddings, Reactome pathways, Gene Ontology biological processes, CORUM protein complexes, CollecTRI regulatory relationships, Human Protein Atlas annotations, Gene Ontology molecular functions, Gene Ontology cellular components, Human Phenotype Ontology annotations and WikiPathways pathways. The prior representations were combined at the level of gene similarity kernels. Missing annotations contributed zero features for the corresponding view.

For within-context models, within each training partition the continuous DepMap, GenePT and STRING features and log1p-transformed Human Protein Atlas profiles were standardized using the mean and standard deviation of the represented training targets, with standard deviations bounded below by 10^—6^. Feature vectors were then normalized to unit Euclidean norm. For annotation views, binary gene-by-term membership vectors were weighted by the inverse square root of term frequency and normalized to unit Euclidean norm. The view-specific kernel *K*_*v*_ was defined as the inner product of two normalized vectors. Prior kernels were combined uniformly or with optional weighting based on alignment with training responses. For alignment weighting, we first removed the fitted response intercept, excluded every perturbation-target gene in the prior bank from the output-gene axis used for alignment, and normalized each remaining response vector to unit norm. Their Gram matrix defined a response kernel. Both this kernel and each prior kernel on the training conditions were double centered. Alignment *a*_*w*_ was the non-negative part of the cosine similarity between their off-diagonal entries; it was set to zero when either vector had negligible norm. For *W* included prior views, the weights and combined kernel were

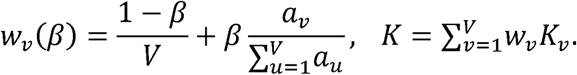

The mixing coefficient *β* ∈ [0,1] controls the alignment contribution. *β* = 0 gives uniform weights and *β* = 1 gives normalized alignment weights. When all alignments were zero, each view received weight 1/*V*. The full combined kernel was retained for within-context regression. Alignment and prior transformations were recomputed from the responses and perturbation targets available to the corresponding fit. The main single- and double-gene benchmarks used all 12 priors with *β* = 0.5.

### Single and double perturbations

Single-perturbation models were fitted separately in each context. Let *Y* contain the training response vectors as rows, *b* be their unweighted mean, and *R* = *Y* — **l***b*. We double centered the training kernel. For a query cross-kernel, we subtracted the training-column means and the query’s mean similarity to training targets, then added the training-grand mean. The predicted response was

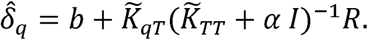

The ridge parameter was selected from 0.01, 0.1 and 1 using validation perturbation-discrimination scores, averaging the L2 and cosine scores across contexts within targets and then across targets. Selected models were refitted on training and validation targets, recomputing feature transformations, response alignment and intercepts. Targets lacking prior coverage received the intercept.

We represented a condition *S* as a set of one or two perturbed genes. For a gene kernel *k*, the shared first-order condition kernel was *K*_*s*_*(S, T) = ∑*_*g*∈*s*_*∑_h_*_∈*T*_*k(g, h)*. Let *l*_*2*_*(S)* indicate that *S* contains two genes. A pair-specific calibration kernel was *K*_*p*_*(S, T) = l*_*2*_*(S)l*_*2*_*(T)K*_*s*_*(S, T)*. For two pairs, the symmetric interaction kernel was

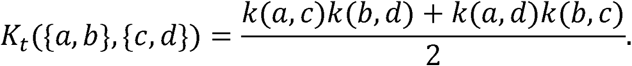

This interaction kernel was zero when either condition was a singleton. The complete condition kernel was *K* = *K*_*s*_ + λ_*p*_*K*_*p*_ + λ_*t*_*K*_*t*_. For response alignment, we first lifted each prior view to this condition kernel using the selected structural coefficients. Its centered off-diagonal entries were compared with the response kernel defined above. The resulting weights were applied to the gene kernels before constructing the final condition kernel, retaining cross-view products in the interaction term.

Single and double conditions were fitted jointly, with equal total weight assigned to each represented perturbation arity. Specifically, condition *i* received weight *W*_*ii*_ = *n*/(*L n_a_i__*), where *n* is the number of training conditions, *a*_*i*_ is its arity, *L* is the number of represented arities and *n*_*a_i_*_ counts conditions of that arity. The intercept design contained the arity *a*_*i*_ and, when enabled, an indicator for double perturbations. The weighted intercept fit was removed before constructing the response kernel for alignment. Kernel coefficients *A* and unpenalized intercept coefficients *B* were estimated simultaneously for all output genes by

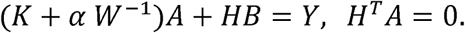

For a query, the predicted response was *K*_*qT*_*A + H*_*q*_*B*, followed by addition of the training-control mean and non-negative expression projection. Double-perturbation observations therefore contributed to estimation of the shared single-gene component. The pair-only kernels and intercept were omitted when no training combinations were available, yielding an additive response before expression projection.

Structural coefficients and the ridge penalty were selected in two stages using seen-0 validation data. Structural coefficients were selected by validation MSE. The full-response, 12-prior model then selected its ridge penalty from 0.01, 0.1 and 1 using mean L2 and cosine perturbation-discrimination scores. Parameters were shared across seen 0, seen 1 and seen 2, with alignment fitted separately to the available training responses. Validation pairs were retained for selection only.

### Prior ablations and alignment weighting

To test gene–prior correspondence, we reassigned gene identities in the prior kernels while retaining the measured perturbation responses and the original training, validation and test partitions. After training-specific feature preprocessing, the same gene-label permutation was applied to the rows and columns of all 12 view kernels. Permutations were restricted to groups of genes with identical coverage signatures across views, contexts and the fitting and refitting stages. Genes in groups of at least two members were reassigned without fixed points; singleton groups remained unchanged. This procedure preserved each target’s prior coverage and each view kernel’s eigenvalues over the full gene universe. We evaluated ten joint permutations, recomputing response alignment and selecting the ridge penalty for each permutation. Control scores were averaged across the ten fitted permutation models (Extended Data Fig. 1a,b).

Prior-number analyses used two random cyclic orders of the 12 views. For each subset size of 2, 4, 6, 8 or 10 priors, the 12 cyclic rotations of each order generated 24 distinct subsets, with every view occurring equally often. All 12 single-view models, all 12 leave-one-view-out models and the full model supplied the results for 1, 11 and 12 priors, respectively. Prediction scores were averaged across subsets at each size; expression predictions were not ensembled. The interquartile range described variability across prior compositions (Extended Data Fig. 1c,d).

Individual prior contributions were assessed by removing each view in turn. We also compared the full model with a three-prior model comprising DepMap, GenePT and STRING, and with a four-prior model that additionally included the log1p-transformed Human Protein Atlas profiles. Each configuration used *β* = 0.5, with weights normalized over the included views, and selected its ridge penalty from 0.01, 0.1 and 1 using the validation procedure described above. These analyses used the five single-gene datasets and their three computational splits (Extended Data Figs. 2 and 3a).

For training-proximity stratification, each held-out target was assigned its maximum similarity to a fitting target under the equally weighted, unpermuted prior kernel. Similarities were averaged over the relevant contexts and computational splits, and targets were divided into low-, intermediate-and high-proximity tertiles within each dataset. The full model and the mean joint-permutation control were compared within each tertile, with datasets weighted equally (Extended Data Fig. 3b).

For double perturbations, CORUM was removed while retaining the structural coefficients of the full model. The remaining view weights were recomputed with *β* = 0.5, and the ridge penalty was selected from 0.01, 0.1 and 1 using seen-0 validation data. That penalty was shared across seen 0, seen 1 and seen 2, with each model fitted to the responses available in the corresponding setting. The comparison used four datasets and five folds per dataset (Extended Data Fig. 5).

Alignment weighting at *β* = 0.5 and uniform weighting at *β* = 0 were compared by rescoring their original predictions on the same supported test conditions, output-gene axis and control references, using the common scoring procedure described below. Each model’s ridge penalty had been selected using the same validation criterion and candidate set. For the sensitivity analysis, we fitted the full 12-prior single-gene model separately at *β* = 0,0.25,0.5,0.75,1, selecting the ridge penalty independently for each value. Mean cosine PDS and DEG AUPRC were summarized with equal weights for the five datasets. *β* = 0.5 remained the configuration used in the main benchmarks (Extended Data Fig. 4).

### Source uncertainty for transfer across cell lines

For prediction across cell lines, the 12 prior kernels were equally weighted over a fixed universe of 2,023 perturbation targets. We retained the leading 512 eigencomponents of the combined kernel as the representation *φ_g_*. The discarded diagonal was retained as *r_g_* = *K*(*g, g*) — ‖*φ_g_*‖^2^, with negative roundoff values set to zero. Augmenting the representation with an intercept gave *z*_*g*_ = [1, *φ_g_*] and the gene covariance

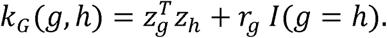

A Gaussian process (GP) ^27^ was fitted independently for each source cell line to estimate the relative uncertainty of its measured responses. For a source condition, the sampling-noise variance was the mean across output genes of the sum of the perturbation and control sample-mean variances. This scalar was divided by the source response scale, defined as the mean across genes of the population variance of condition-level responses, with a floor of 10^—12^ on that scale. The normalized noise was bounded below by 10^—6^. The covariance was evaluated as a low-rank plus diagonal matrix, using a Cholesky factorization of the corresponding feature-space system.

For a query perturbation *g* and source cell line *s*, let *T*_*s*_ denote the measured training perturbations and *E*_*s*_ be the diagonal matrix of normalized sampling-noise variances. All covariance blocks were evaluated using *k*_*G*_. We first computed the latent GP posterior variance on the source-normalized scale,

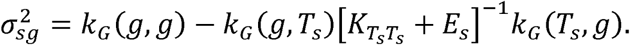

We then defined the relative posterior variance as the dimensionless posterior-to-prior variance ratio,

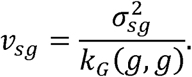

This ratio was used to weight source contributions. The source point estimate was the measured response *δ_sg_*. Only sources in which the query had been measured contributed to the transferred prediction. The prior representation was shared across sources, whereas response scales, sampling variances and posterior covariance factors were fitted separately for each source.

### Optimal transport

We sampled up to 512 control cells per cell line without replacement. Controls were centered within each cell line. We obtained a common 128-dimensional basis *U* by exact eigendecomposition of the Gram matrix of the stacked controls, weighting each cell line equally. Sample covariances in this basis used denominator *n* — 1. Covariance eigenvalues were bounded below by the larger of 10^—8^ times their mean and 10^—12^.

Let *S*_*s*_ and *S*_*t*_ be the source and target control covariances in this basis. The Gaussian optimal transport map ^28^ was

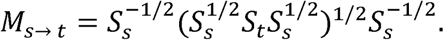

We parameterized the transported response by a transport strength *η*: *δ_sg_* + *ηδ_sg_U*(*M_s → t_* — *l*)*U^T^*. This construction transforms the component in the control-derived basis and preserves the orthogonal component. The transferred response used the full map, with zero covariance shrinkage. The main benchmark used *η* = 1.

Control-cell similarity used the squared difference between cell-line control means, standardized gene-wise by the source-averaged control variance and averaged over genes. Variances were bounded below by 5% of their across-gene mean. Denote this distance by *d*_*st*_ and the median distance between source cell lines by *ℓ*, bounded below by 10^—12^. For the measured-source set *S*(*g*), weights were

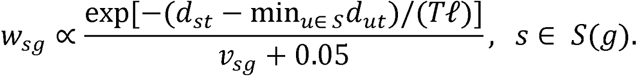

Source weights were normalized over sources containing the query perturbation. The temperature was selected by leaving out source cell lines in turn and maximizing control-similarity-weighted discrimination, subject to an error constraint relative to the fixed-temperature baseline.

### Data and benchmarks

For single-gene perturbation, we used the Tian CRISPRi and CRISPRa^30^, Adamson^6^, Norman single-perturbation^31^ and Frangieh^32^ datasets from scPerturb^33^ (Zenodo record 13350497); for double-gene perturbation, we used Norman^31^, Replogle exp6^8^, Wessels lane L4^34^ and Dixit K562 high-MOI^5^ data; for perturbations across cell lines, we used the released Replogle–Nadig essential-gene benchmark^9,18^ and unseen-context splits from PIE^35^.

Single-perturbation comparisons included biolord^36^, Scouter^13^, TxPert^12^, GEARS^3^, STAMP–Geneformer^37,38^ and RegFormer–GEARS^3,39^. All used the same target partitions, output axes and control references. Comparator learning rates and stopping points were selected using validation expression RMSE, averaging contexts within targets and then targets. Models were refitted on training and validation data for the selected number of optimizer updates. Double-perturbation comparisons included GEARS, Scouter, TxPert, STAMP and GPerturb^40^, together with no-change and perturbed-centroid predictors.

Cross-cell-line STATE^4^, STACK^41^ and TxPert models were trained separately for each held-out cell line using the benchmark’s allowed training and validation pairs. STATE and TxPert started from random initialization; STACK used local masked-gene pretraining followed by in-context posttraining. STATE used cell sets of length 64, eight transformer layers and an energy-distance loss. STACK used nine layers and 128-cell sets, with measured same-perturbation source cells as prompts. TxPert used its cross-cell-line graph-attention model adapted to the 6,642-gene output axis.

### Perturbation metrics

Expression MSE and Pearson correlation were calculated between predicted and observed mean profiles or their expression changes relative to the control mean. For shared-response Pearson correlation, both predicted and observed profiles were centered on the mean of all retained training perturbation cells in the same context. This reference weighted perturbations by their retained cell counts. For top-100 MSE, Pearson correlation and sign agreement, genes were ordered by the absolute observed Welch t statistic, using a stable order for ties. Observed differentially expressed genes (DEGs) required a two-sided Welch test with Benjamini–Hochberg-adjusted P < 0.05 and an absolute mean log1p expression change > 0.1. Predicted changes were ranked by absolute magnitude for precision–recall and top-100 recovery. For perturbation discrimination, each predicted response was compared with observed responses for common-supported test targets in the same dataset, context and split. The query’s directly targeted gene was removed from every comparison when present on the output axis. With *N* candidates, *b* strictly closer candidates and *t* other candidates tied within 10^—12^ of the correct response, the score was 1 — (*b* + *t*/2)/(*N* — 1). Distances were squared Euclidean or cosine. Predicted responses with root-mean-square magnitude at most 10^—7^ were set to zero; zero-vector cosine similarities were zero. Pools with fewer than two candidates were undefined. Scores were averaged over contexts within each target and split, over splits within each target, and finally over unique targets. Dataset summaries weighted targets equally, and radar plots weighted datasets equally.

For each test pair and context, the top 20 genes were those with the largest absolute observed change relative to the independent scoring-control mean. The opposite-direction fraction counted genes for which the product of predicted and observed changes was negative; a zero change was not counted as an opposite direction. Normalized MSE was the ratio of aggregated prediction MSE to aggregated squared observed change on the same top-20 genes. Contexts were averaged within pair, pairs within split and neural-model seeds within split. Across-dataset summaries weighted the four datasets equally. For combination identification, each observed response was compared with predictions for all held-out pairs in the same context and split by Pearson correlation across the 5,000 output genes. Zero-norm correlations were assigned similarity zero. If *b* predictions had higher similarity and *t* tied predictions included the correct pair, fractional top-*K* credit was max[0, min(1, (*K* — *b*)/*t*)]. Ties used tolerance 10^—12^. Repeated scores were averaged within pair before averaging across distinct pairs. Expression examples and residual displays used fixed dataset, split and availability settings. Observed and predicted departures from additivity were calculated with the same measured singleton means and scoring-control reference.

Response-direction agreement was the fraction of observed DEGs with matching signs of observed and predicted log-fold change. Conditions with no observed DEGs received zero and remained in the average. L1 discrimination ranked the observed response among all test conditions in the same target cell line after removing the query target gene. With zero-based ordinal rank *r* and *N* candidates, the score was 1 — *r/N*. Candidate conditions were ordered by perturbation name, and distances were ranked with NumPy argsort. Pearson correlation compared the 6,642-dimensional predicted and observed response vectors. Spearman correlation compared log-fold changes only among observed DEGs, using average ranks for ties. Constant-vector correlations and Spearman correlations with fewer than two genes were undefined and omitted.

For DEG overlap, let *N*_*DE*_ be the number of observed DEGs. Predicted significant genes were ranked by absolute log-fold change, with stable tie ordering, and the overlap of the first *N*_*DE*_ genes with the observed DEG set was divided by *N*_*DE*_. All predicted significant genes were used when fewer than this number were available. Conditions with no observed or predicted DEGs received zero. The direction, Pearson, Spearman and overlap metrics retained the directly targeted gene. DEG-count plots included all test conditions at adjusted P below 0.05.

### Cardiac reprogramming data

We used day-14 expression measurements and perturbation and sample-batch annotations from the in vivo cardiac reprogramming Perturb-seq study by Cai et al. (PMID: 41916284; Zenodo record 19185135). ^29^ The analysis used the supplied normalized expression scale and four sample batches. The cardiac score was the mean expression of *Tpm1*, *Tnnt2*, *Ryr2*, *Ttn*, *Ctnna3*, *Myl4*, *Myh6* and *Actc1*.

For each of 20 cell splits, all query groups with at least 15 cells contributed five input cells and reserved all remaining cells as a disjoint reference. Other targets with at least five cells in the same batch contributed all their cells to fitting. This yielded 77 target-by-batch cases representing 33 targets. The ten-cell sensitivity analysis required at least 20 cells and included 53 cases representing 24 targets. Non-targeting controls were divided into disjoint input and reference groups within each batch and split. The same allocations were used for all estimators.

We evaluated the single-perturbation kernel-ridge model at the identities of these measured input targets. Each fit used input mean expression profiles, a training-target intercept and the 12-prior aligned kernel; negative fitted means were set to zero. Response alignment excluded assayed target genes and the additional symbols *Becn1*, *Ifnar2* and *Wwtr1*, leaving 25,804 of 25,944 genes. Five cell folds within each target were used to select one ridge parameter for each batch, input budget and outer split from 0.01, 0.1 and 1 by cardiac-score mean squared error (MSE), weighting targets and folds equally. Alignment and fitting were repeated within each fold. Ties within 10□¹² favored the larger parameter. Reference cells and reference controls were used only for evaluation.

KiPERT and input sample means were compared with reference means for mean cardiac-gene expression before control subtraction and for expression changes. Predicted changes used input controls, whereas reference changes used reference controls. Squared errors were averaged over cell splits within each target-by-batch case, over eligible batches within each target, and equally over targets. Relative error reduction was one minus the ratio of KiPERT MSE to sample-mean MSE. Paired 95% percentile intervals used 2,000 resamples of targets, retaining their batches, splits and predictions. The resampling unit was the perturbation target, with repeated cell splits nested within targets.

### Transfer of *Calr* response across batches

For each day-14 target batch, KiPERT transferred measured responses of the same perturbation from the other batches. Eligibility required at least five target-batch cells and at least two source batches with five cells each, yielding 211 target-by-batch queries. Target-batch perturbed cells were withheld from fitting and prediction, while its controls defined the transport geometry. The Gaussian control-transport equation above was used with confidence weights proportional to 1/(v + 0.05) and a uniform context-similarity factor. Confidence was estimated separately from each source’s observed responses and sampling variances using the full, equally weighted 12-prior kernel plus intercept. Transported changes were evaluated directly on the supplied expression scale.

*Calr* was examined as a previously reported positive reference from the original study. Its complete eight-gene response was evaluated in all four batches, containing 14, 13, 12 and 13 *Calr* cells and 27, 59, 26 and 30 non-targeting controls. Measurement intervals used 2,000 independent cell resamples of *Calr* and control cells within each batch, preserving the gene vectors. Percentile limits were calculated for each gene and for the eight-gene score. These intervals quantify within-batch measurement variation.

---

**Extended Data Fig. 1.**
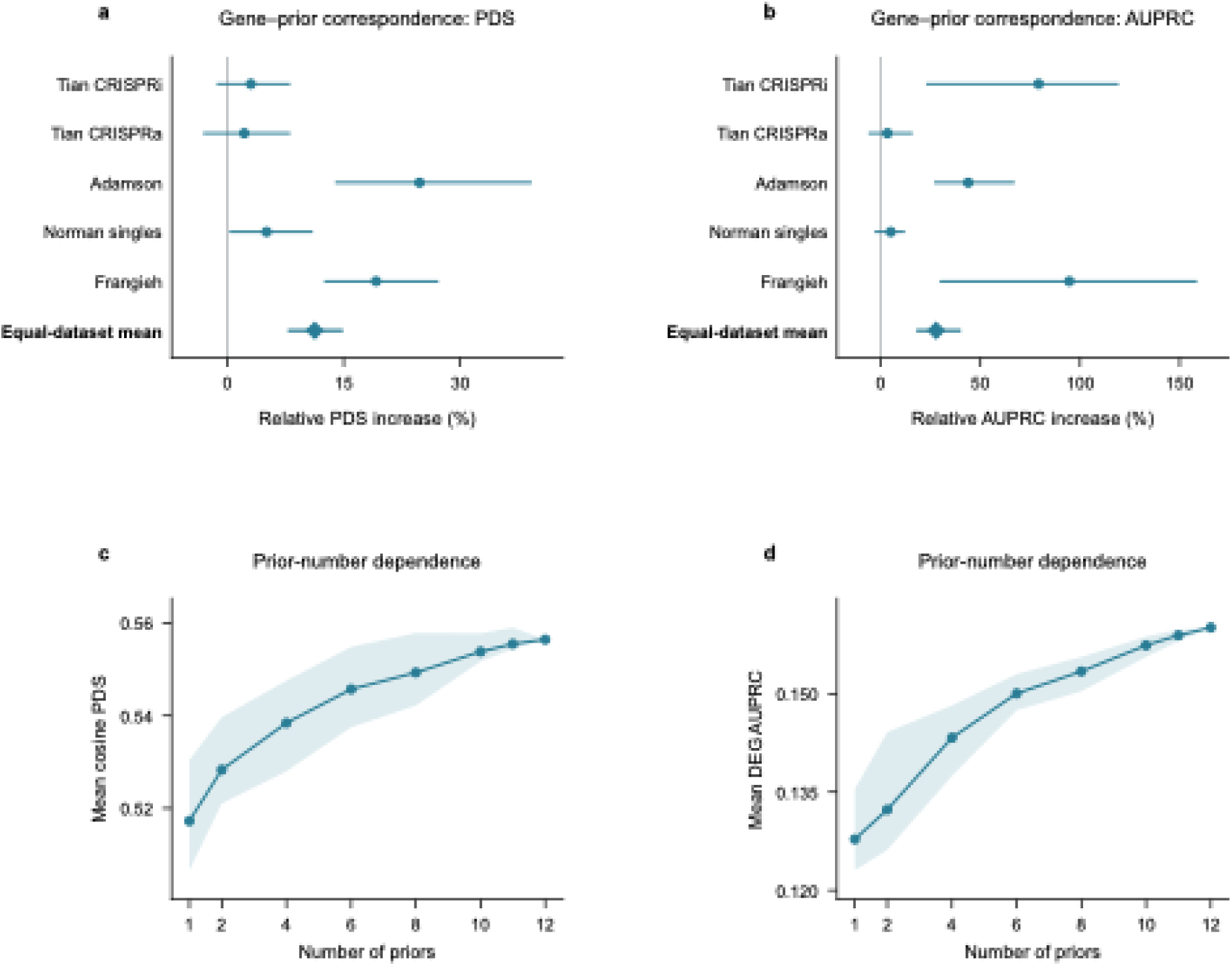
Gene–prior correspondence. **a,b,** Relative increases in cosine perturbation discrimination score (PDS; a) and area under the precision–recall curve for differentially expressed genes (DEG AUPRC; b) for the full-prior model over the mean of ten joint gene-label permutations. The permutations preserve target coverage and each full-universe kernel spectrum. Circles denote dataset effects and diamonds denote effects calculated from equally weighted dataset means. Horizontal lines show pointwise 95% paired target-cluster bootstrap intervals. Relative increases are 100 × (full-model score/control score − 1). **c,d,** Equally weighted mean PDS (c) and DEG AUPRC (d) across prior-set sizes. Size 12 is the full model. Shading denotes composition quartiles.

**Extended Data Fig. 2.**
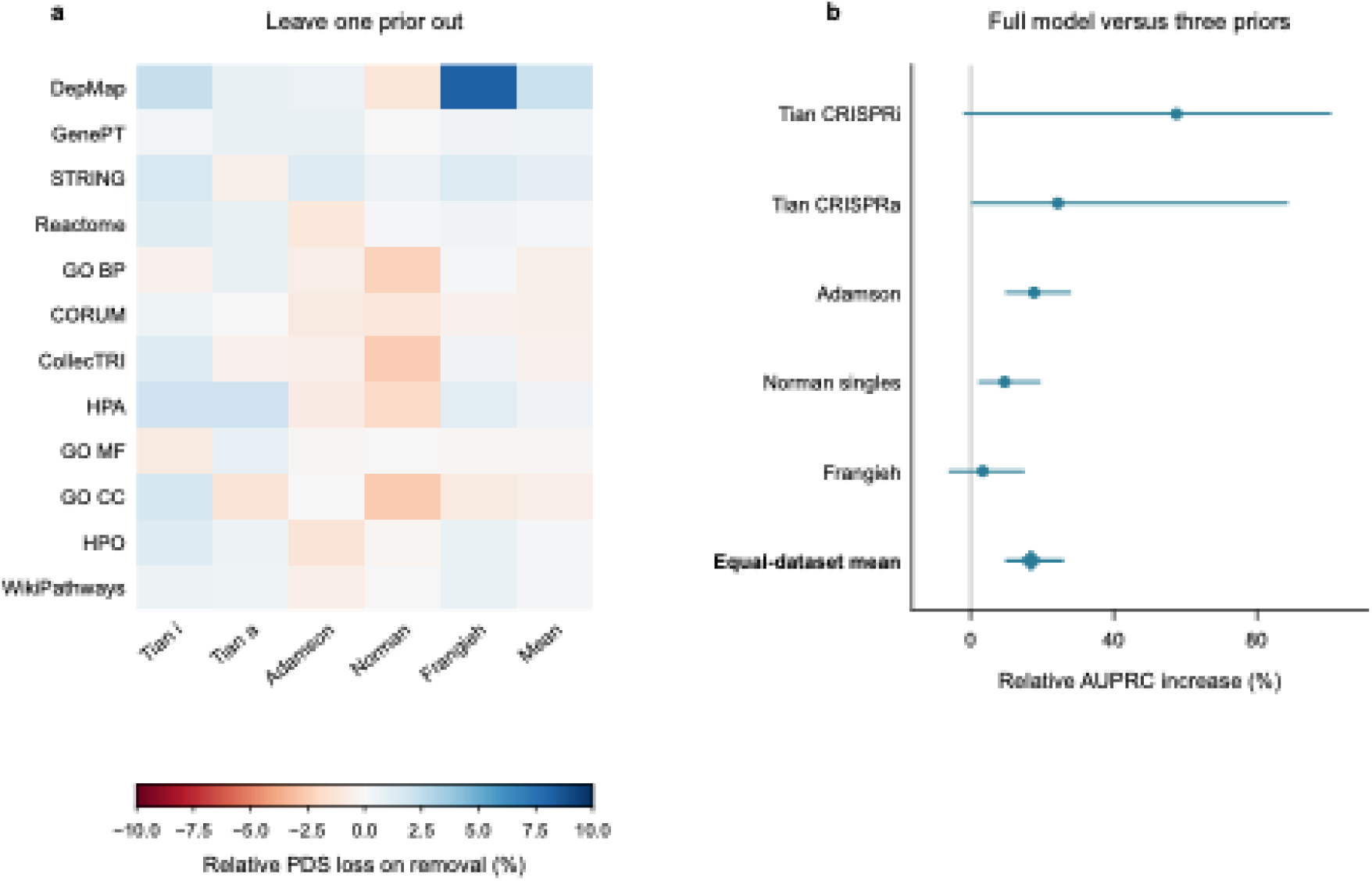
Contributions of individual priors. **a,** Relative loss in cosine perturbation discrimination score (PDS) after removal of each of the 12 priors, calculated as 100 × (full-model PDS − ablated-model PDS)/full-model PDS. Blue denotes lower PDS after removal and red denotes higher PDS. The Mean column uses equally weighted dataset scores. **b,** Relative increase in area under the precision–recall curve for differentially expressed genes (DEG AUPRC) for the full model over the DepMap/GenePT/STRING model, calculated as 100 × (full-model AUPRC/three-prior AUPRC − 1). Circles denote dataset effects, the diamond denotes the effect calculated from equally weighted dataset means, and horizontal lines denote pointwise 95% paired target-cluster bootstrap intervals from 2,000 resamples.

**Extended Data Fig. 3.**
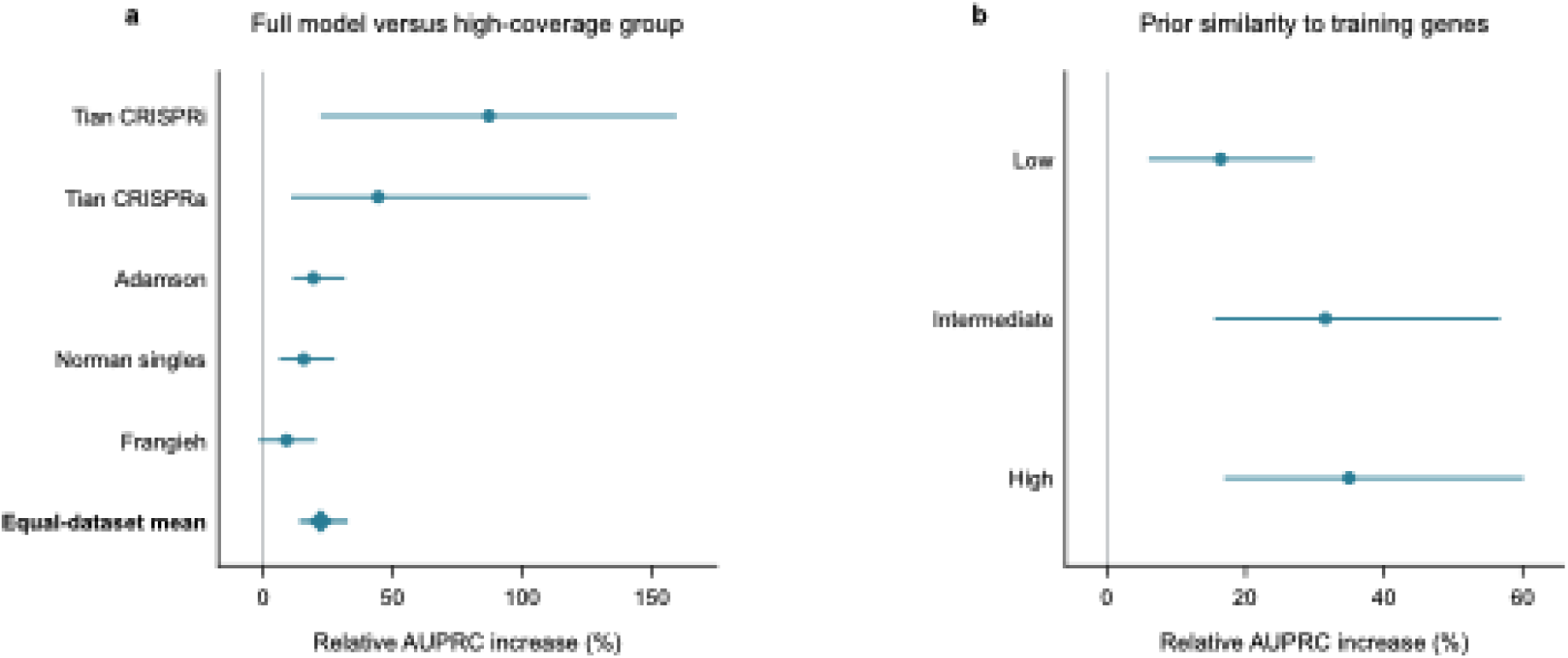
DEG recovery analysis. **a,** Relative increase in area under the precision–recall curve for differentially expressed genes (DEG AUPRC) for the full model over the high-coverage group comprising DepMap, GenePT, STRING and the Human Protein Atlas. Circles denote dataset effects and the diamond denotes the effect calculated from equally weighted dataset means. **b,** Relative DEG AUPRC increase over the mean joint-permutation control in low-, intermediate- and high-proximity groups. Proximity is nearest-training-target similarity under the uniform original kernel, averaged across contexts and computational partitions before defining within-dataset tertiles. Relative increases are 100 × (full-model score/reference score − 1). Horizontal lines show pointwise 95% percentile intervals from 2,000 paired target-cluster bootstrap resamples.

**Extended Data Fig. 4.**
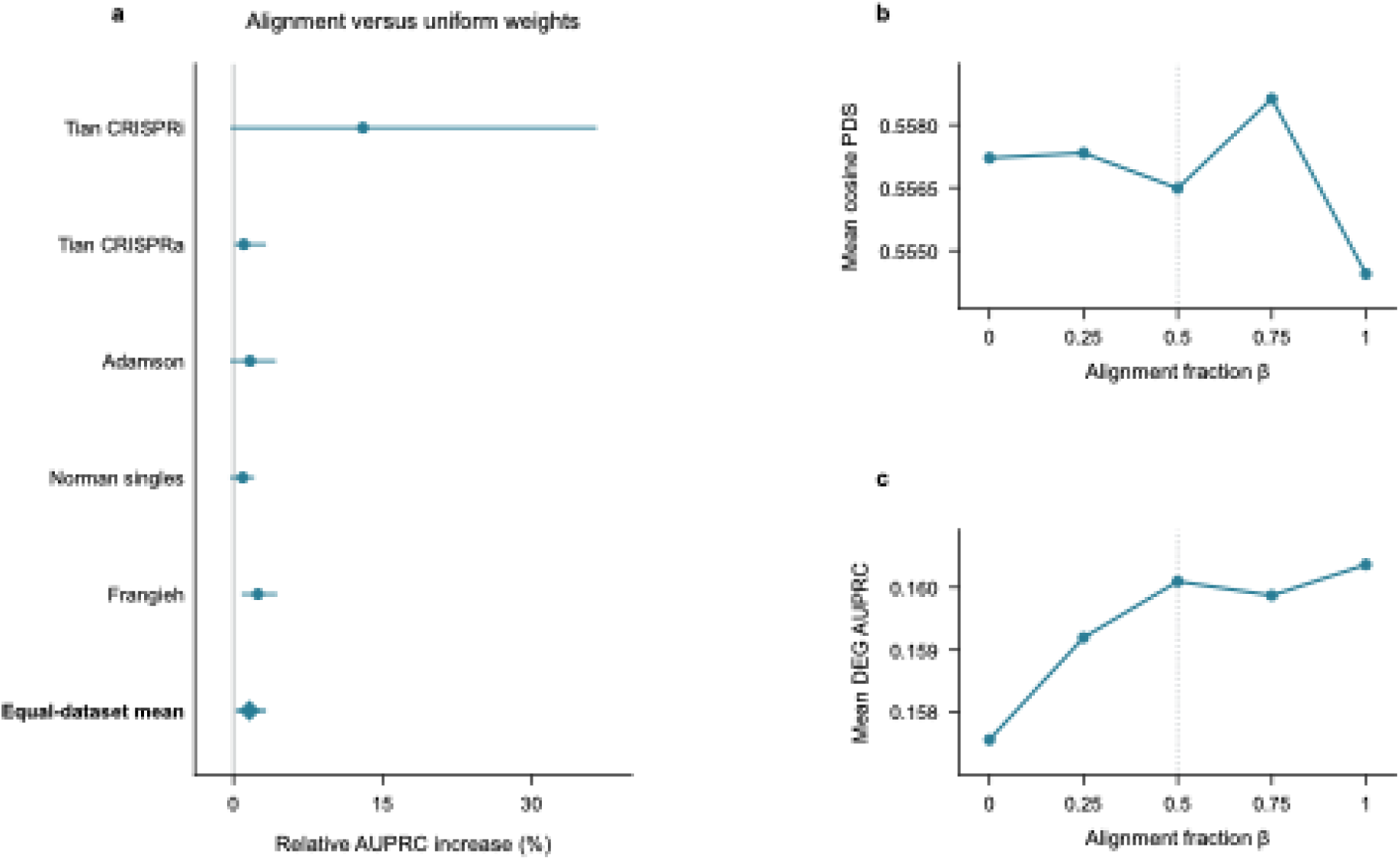
Optional alignment weighting and sensitivity analysis. **a,** Relative change in area under the precision–recall curve for differentially expressed genes (DEG AUPRC) between alignment weighting (*β* = 0.5) and uniform weighting (*β* = 0). Relative change is 100 × (aligned AUPRC/uniform AUPRC − 1). Circles denote dataset effects, the diamond denotes the effect calculated from equally weighted dataset means, and horizontal lines denote pointwise 95% percentile intervals from 2,000 paired target-cluster bootstrap resamples. **b,c,** Equally weighted mean cosine perturbation discrimination score (PDS; b) and DEG AUPRC (c) at the five alignment fractions. The dotted line marks *β* = 0.5, used in the main benchmarks.

**Extended Data Fig. 5.**
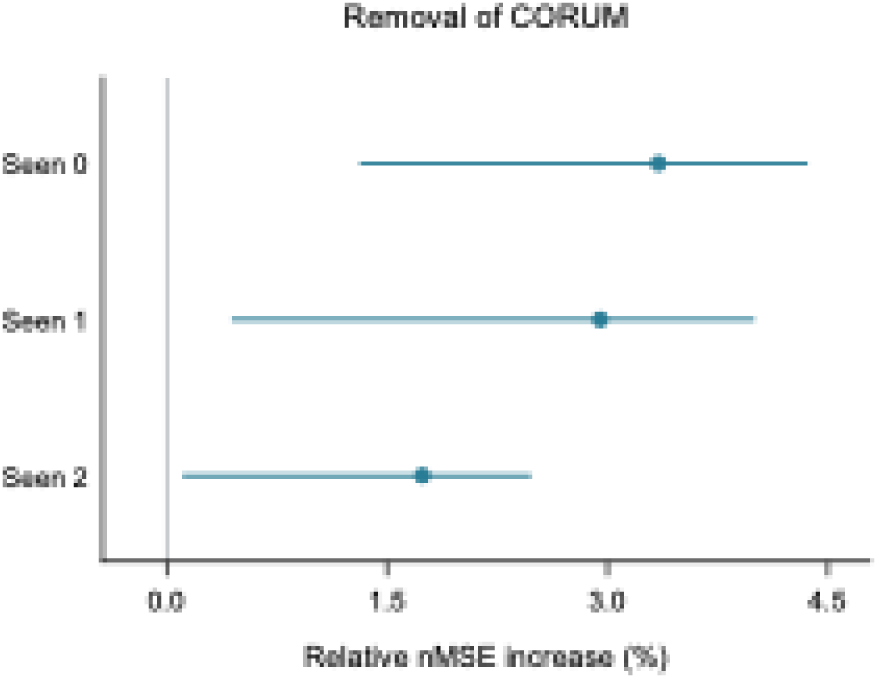
Contribution of CORUM to double-perturbation prediction. Relative increase in top-20 normalized mean squared error (nMSE) after removing CORUM, calculated as 100 × (nMSE without CORUM/full-model nMSE − 1). Positive values indicate lower error for the full model. Horizontal lines show pointwise 95% percentile intervals from 2,000 paired gene-pair-cluster bootstrap resamples.

**Extended Data Fig. 6.**
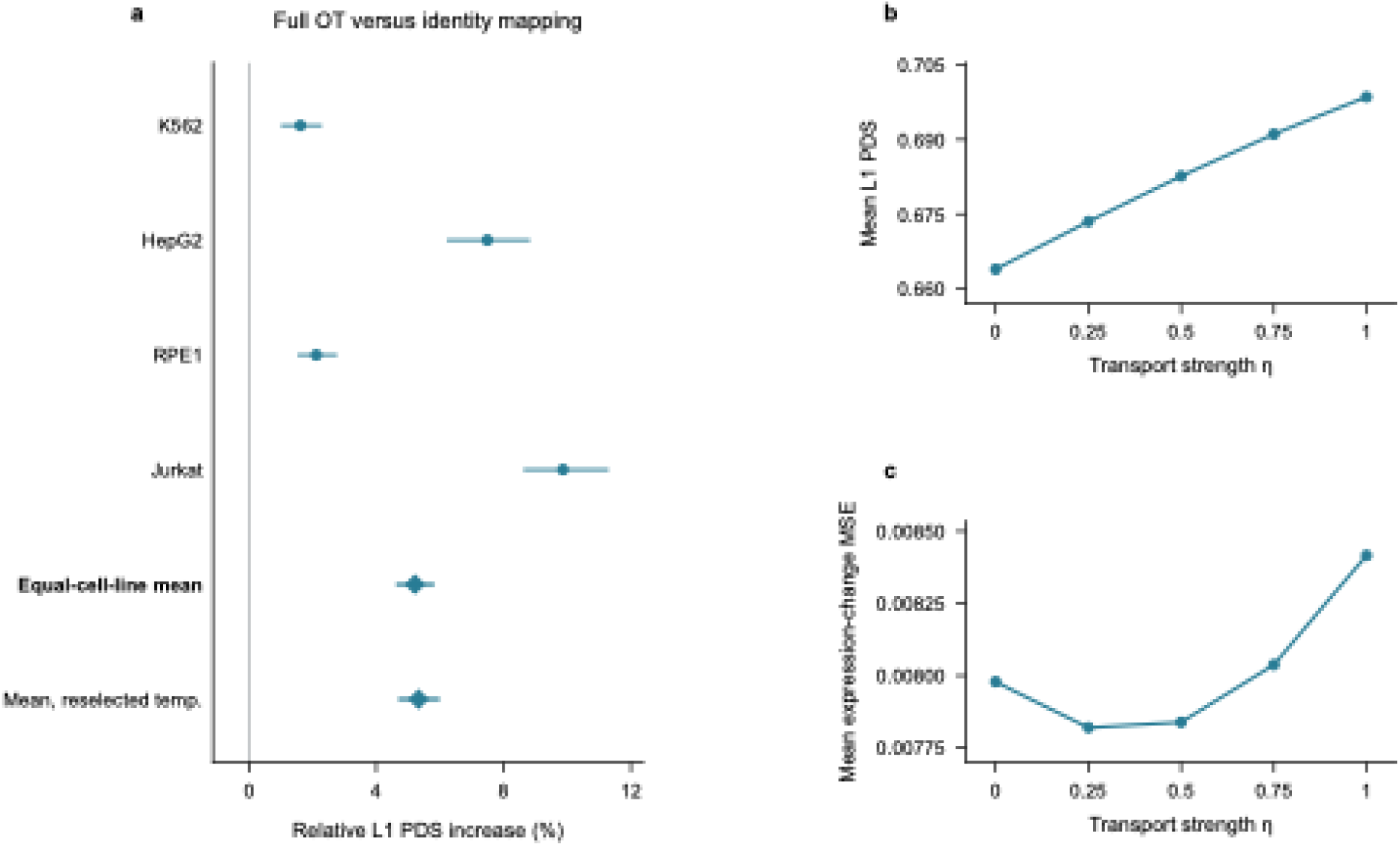
Optimal transport and sensitivity to transport strength. **a,** Relative increase in L1 perturbation discrimination score (PDS) with full optimal transport (OT) compared with identity mapping, calculated as 100 × (OT PDS/identity PDS − 1). **b,c,** Mean L1 PDS (b) and expression-change mean squared error (MSE; c) versus transport strength *η*, with source weights. *η* = 0 gives identity mapping and *η* = 1 gives full OT.

## Notes

### Competing Interest Statement

The authors have declared no competing interest.

